# A Hybrid Multiscale Model for Predicting CAR-T Therapy Outcomes in Solid Tumors

**DOI:** 10.1101/2025.11.17.688869

**Authors:** Mohammad R. Nikmaneshi, Lance L. Munn

**Affiliations:** Edwin L. Steele Laboratories, Department of Radiation Oncology, Harvard Medical School and Massachusetts General Hospital, Boston, Massachusetts, 02114, United States of America

## Abstract

T cell distribution within tumors (“tumor hotness”) critically determines immunotherapy success. However, despite numerous strategies to enhance intratumoral T cell accumulation—such as multi-target CAR-Ts and combinatorial approaches—limited mechanistic understanding of T cell–microenvironment interactions has constrained progress. To address this, we developed a physiological mechanistic model of the 3D tumor microenvironment (TME) to evaluate CAR-T performance under environmental fluctuations and different infusion strategies. The model integrates key vascular (rolling, adhesion, endothelial suppression) and interstitial (ECM density, metabolic competition, chemokine sensitivity) barriers. Our simulations reveal that collagen density and metabolic competition dominate CAR-T efficacy. Enhancing vascular adhesion improves infiltration but remains limited by collagen and metabolism. Endothelial suppression markedly reduces tumor hotness, while its alleviation enhances response. Systemic infusion yields higher tumor hotness than intratumoral delivery, but combined routes or reduced collagen restore efficacy even in dense tumors. This mechanistic framework enables rational optimization of CAR-T strategies.

**Significance Statement:** The success of immunotherapies such as CAR-T cells depends on their ability to infiltrate and persist within solid tumors, yet the mechanisms that govern this process remain poorly understood. Using a mechanistic 3D model of the tumor microenvironment, we quantitatively dissected how vascular and interstitial barriers—including endothelial suppression, collagen density, metabolic competition, and chemokine cues—shape CAR-T distribution (“tumor hotness”). Our results reveal that stromal and metabolic constraints, rather than vascular adhesion alone, dominate CAR-T efficacy. This framework bridges molecular, cellular, and tissue-scale mechanisms, providing a quantitative foundation for optimizing CAR-T design and delivery strategies to overcome resistance in solid tumors.

## Introduction

The tumor microenvironment (TME) is a dynamic ecosystem where cancer, immune, and stromal cells interact within a structurally and physiologically abnormal milieu. Effector T cells—critical mediators of anti-tumor immunity—must infiltrate this environment to exert cytotoxic effects. The extent of T cell accumulation within tumors, often referred to as tumor “hotness,” is a key determinant of immune checkpoint blockade (ICB) and other immunotherapy outcomes ^1,2^. Tumors with dense T cell infiltration are considered immunologically “hot,” whereas poorly infiltrated tumors are “cold.” Understanding the mechanisms governing T cell infiltration and distribution is therefore essential for optimizing therapeutic efficacy.

Tumor vasculature and spatial heterogeneity present major barriers to T cell entry. Tumor vessels frequently exhibit irregular architecture, poor perfusion, and low expression of adhesion molecules ^3–5^, which reduce T cell–endothelial interactions. T cells that encounter the vessel wall must complete a multistep adhesion cascade—rolling, integrin activation, and firm adhesion—before transmigration ^6–8^. However, tumor-associated endothelial cells often display immunosuppressive phenotypes, expressing inhibitory molecules such as PD-L1 or FasL ^9,10^ and secreting factors such as TGF-β that interfere with adhesion, arrest, and activation ^11–16^. These mechanisms act as vascular checkpoints that suppress or eliminate T cells before entry into tumor tissue. Even after extravasation, T cells encounter a dense extracellular matrix, limited nutrients, and disorganized chemokine gradients ^17,18^, all of which hinder migration and effector function. Together, these biophysical and biochemical barriers limit both the number and efficacy of tumor-infiltrating lymphocytes.

Mathematical modeling provides a quantitative framework to dissect these multiscale interactions and predict how microenvironmental factors influence T cell distribution. Temporal (non-spatial) models, often based on ordinary differential equations (ODEs), describe immune–tumor population dynamics ^19–29^. While valuable for fitting clinical data and predicting systemic behavior, they overlook spatial heterogeneity and local interactions critical to the TME. Spatiotemporal continuous models simulate 2D or simplified 3D environments with continuous variables (e.g., oxygen, chemokines, or flow) ^30–36^, capturing spatial gradients but not individual cell variability. In contrast, agent-based models (ABMs) represent individual cells as discrete agents with stochastic rules ^37–41^, enabling simulation of heterogeneous and localized interactions, yet lacking explicit continuous fields. Hybrid discrete–continuous (HDC) models bridge these frameworks by coupling ABMs with continuous fields of signaling molecules, mechanical stresses, and interstitial flow ^42–52^. These 3D hybrid models best represent the spatial and temporal complexity of the TME, integrating stochastic cell dynamics with deterministic microenvironmental processes ^48,49,53^.

Most prior models have examined either systemic immune circulation ^19–22^ or local tumor–immune interactions ^46,47^, leaving a gap in understanding how T cells traverse the vasculature and infiltrate solid tumors within a realistic 3D context. To address this, we developed a hybrid, multiscale 3D model combining agent-based representations of immune and cancer cells with continuous descriptions of flow and signaling in a vascularized TME. The model incorporates key vascular and interstitial barriers—rolling, adhesion, endothelial suppression, extracellular matrix density, metabolic competition, and chemotactic signaling—to quantify their collective impact on CAR-T distribution and efficacy. Built upon a mechanistic hybrid framework and refined through machine learning–based validation, this physiologically grounded model provides a predictive platform for investigating tumor “hotness” and optimizing CAR-T therapy in solid tumors. Details of the model development—from the construction of the generic framework to its conversion into a physiological model describing CAR-T cell interactions with solid tumors—are provided in the Methods section.

## Results

We used the physiological model to evaluate CAR-T performance by varying intravascular and extravascular properties of the TME and using different CAR-T infusion strategies. We first examined CAR-T distribution within the tumor following systemic infusion under varying vascular and interstitial conditions, and then assessed therapeutic outcomes for each scenario. To achieve this, we modeled T cells in two phases: a non-cytotoxic state and a cytotoxic state.

### Intratumor CAR-T distribution is affected by the TME

#### Quantifying intratumor CAR-T distribution

CAR-T cells are infused systemically (intravenously, IV), and their plasma concentration is modeled as a Gaussian function over time (Fig. 1). The spatiotemporal dynamics of CAR-T cells during tumor growth are shown in Fig. 1. After systemic infusion, CAR-T cells enter the tumor microenvironment and distribute both around and within the tumor (violet).

**Figure 1.**
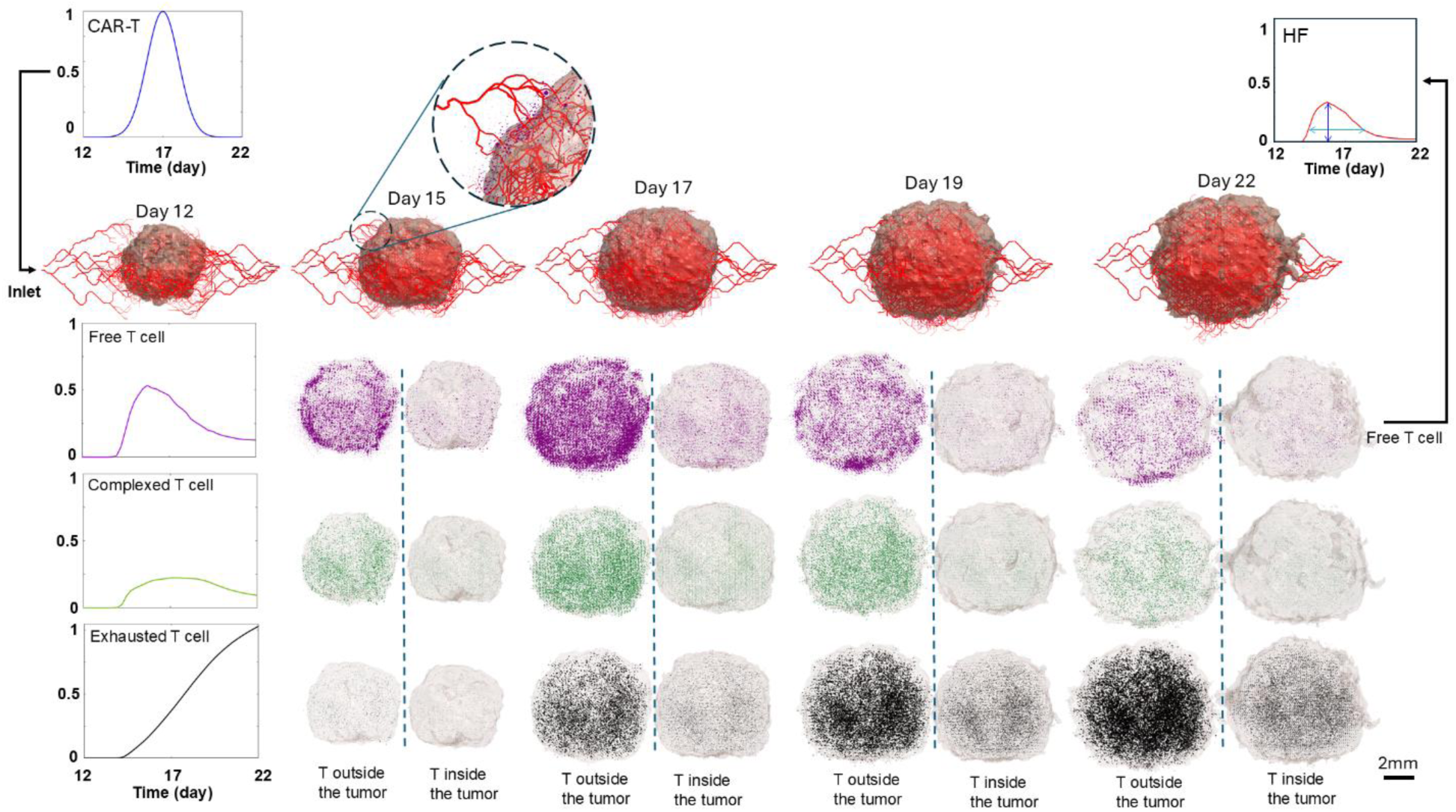
Monitoring T cell trafficking in the tumor microenvironment (TME) following systemic infusion. Spatiotemporal dynamics of T cells within a vascularized tumor are visualized at different time points of tumor growth. Violet indicates free T cells (single cells and colonies), green indicates T cells in complex with cancer cells, and black indicates exhausted T cells engaged with cancer cells. T cell distributions inside and around the tumor are qualitatively compared at each time point. The left panels present quantitative data on the distribution of these T cell populations within the tumor. Based on the spatial distribution of free T cells, the hotness factor (HF) of each infusion is calculated, serving as a metric for evaluating infusion effectiveness in cancer control. In the HF plot, dark and light blue arrows indicate the HF peak and the effective duration corresponding to each HF level.

Infiltrating T cells may form conjugates with cancer cells (green) or become exhausted after prolonged engagement (black). To quantify the extent and uniformity of T-cell distribution within the tumor tissue, we define a *hotness factor* (HF), calculated as the average spatial density of activated CAR-T cells surrounding cancer cells across the entire tumor domain at each time point (Eq. 45, Supplementary Material).

For this calculation, only activated CAR-T cells moving freely within the tumor and not yet conjugated with cancer cells are considered (violet T cells, Fig. 1). We first evaluate HF under a non-cytotoxic assumption to isolate the effects of trafficking and infiltration, since efficient and homogeneous T-cell delivery is a prerequisite for effective therapy. We then incorporate cytotoxic activity using the average killing rate obtained in Fig. 6A of the Methods section to assess how HF modulates tumor control in the following section. The resulting HF of non-cytotoxic free CAR-Ts, represented by the red curve in Fig. 1, reveals that the T cells accumulate primarily in perivascular and peripheral tumor regions, leading to non-uniform intratumoral infiltration and overall low HF values—patterns commonly observed in solid tumors^2,75–77^.

#### Intravascular interactions of CAR-T cells

To explore how cell trafficking mechanisms affect the T cell distribution in tumors, we used the physiological tumor model developed in the Methods section as a baseline and varied parameters related to T cell rolling frequency, firm adhesion intensity, and suppressive factor strength across wide ranges (0.01×, 0.1×, 10×, and 100×). While decreasing rolling and firm adhesion markedly reduced HF by lowering the number of T cells arrested at vessels, increasing their intensity did not appreciably improve HF. This is because, even when T cells are captured and transmigrate, there are barriers to uniform infiltration into the interstitium. The model predicts that firm adhesion is prevalent in small neo-angiogenic vessels with low shear stress, and that rolling adhesion has comparatively less impact on infiltration. Our results further show that, with 100-fold reductions in rolling or firm adhesion relative to baseline, the HF peak is delayed by 0.6 day for reduced rolling and 2.8 days for reduced firm adhesion, while peak HF decreases by 40% and 58%, respectively (Figure 2).

**Figure 2.**
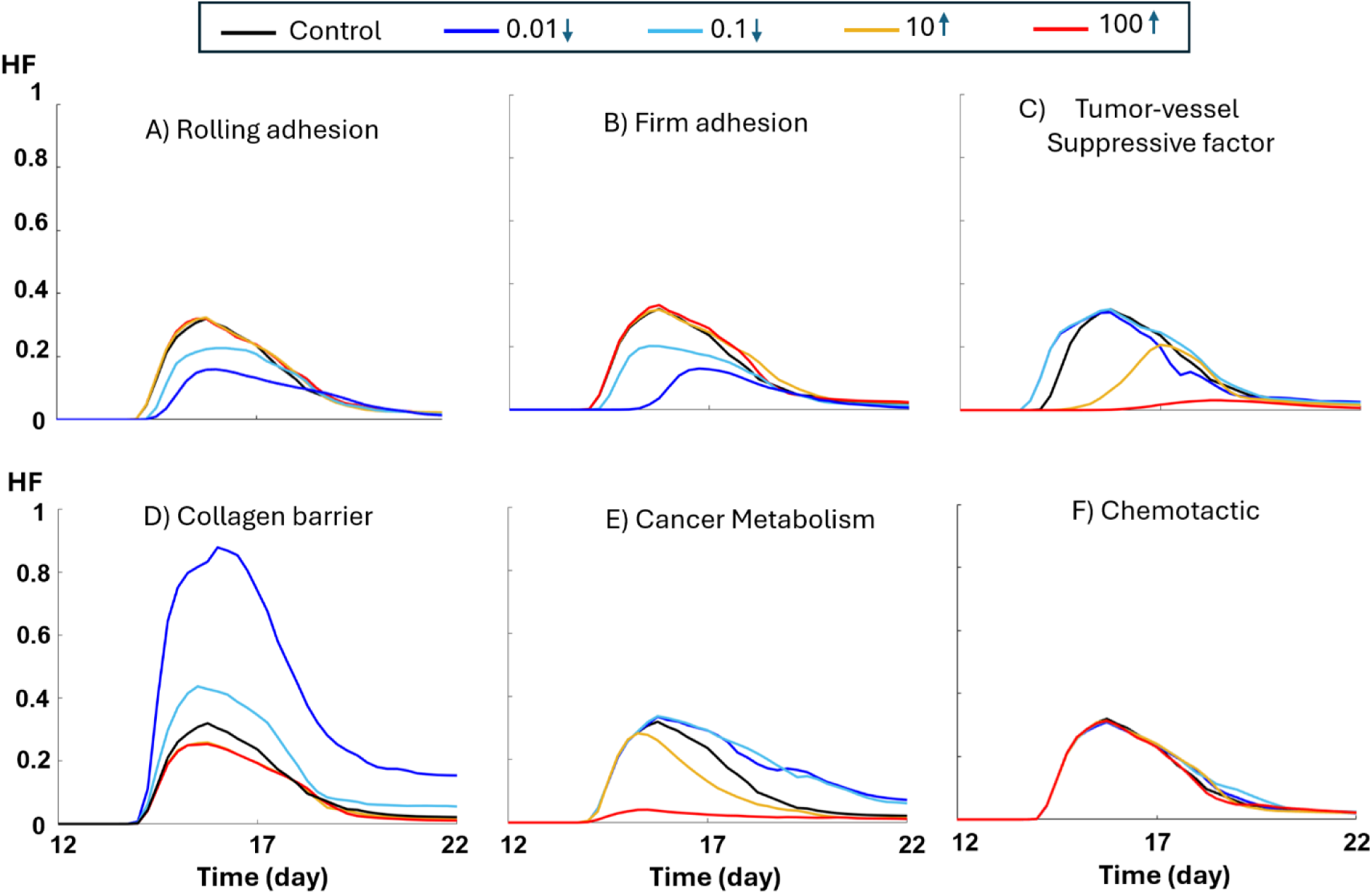
Effects of T cell interactions with vessels, tumor ECM, and tumor. HF profiles are shown following systemic infusion of non-cytotoxic CAR-Ts under distinct mechanisms: (A) rolling adhesion intensity, (B) firm adhesion intensity, (C) tumor-induced SF intensity, (D) tumor-induced collagen concentration; (E) tumor-driven metabolic competition through nutrient consumption and waste accumulation; and (F) tumor-derived T cell–attracting chemokines. The black line represents the baseline control solid tumor, while the tested values (0.01, 0.1, 10, and 100) denote fold changes relative to the baseline parameters for each mechanism.

For suppressive factors, increasing SF intensity strongly decreased HF, ultimately halting T cell infiltration by suppressing T cells before they can exit the tumor vessels. A 10-fold and 100-fold increase in SF reduced HF by 33% and 83%, respectively, and delayed the HF peak by 3.3 days and 8.6 days (Figure 2). In contrast, reducing SF yielded only modest improvements, primarily by slightly expanding the effective duration of HF (Identified by light blue arrow in HF plot in Fig.1) and increasing the chances of early T cell infiltration. Thus, therapies designed to decrease tumor-induced vascular suppression of T cells might protect adherent T cells and allow them to enter the tumor, but their effectiveness remains limited if they don’t penetrate the dense ECM and distribute within the extravascular space. Overall, these mechanistic results highlight the need for reducing SF, enhancing firm adhesion, and possibly increasing rolling adhesion before CAR-T infusion. However, even under optimal conditions, the maximal improvement only achieves a HF of ∼0.4 —still far below the desired value of ∼1, where the entire tumor mass would be uniformly infiltrated by T cells, and each cancer cell would be in close proximity to at least one T cell.

#### Interstitial interactions of CAR-T cells

Our simulations predict that tumor fibrosis and the associated high collagen content play a manor role in determining tumor hotness (Fig. 2D). Reducing collagen significantly increases HF in tumors with the same levels of rolling adhesion, firm adhesion, and SF. Comparison with the intravascular factors in Figs. 2A–C indicates that, in addition to reducing intravascular barriers, T cells must be able to migrate efficiently within the tumor to achieve broader distribution before being suppressed by sequential interactions with multiple cancer cells. Notably, the baseline model, calibrated with a variety of solid tumors, reflects high collagen-induced physical barriers, consistent with experimental observations that dense ECM—particularly collagen—is a hallmark of solid tumors^78^ Consequently, increasing collagen beyond the baseline only slightly reduces HF due to the already high collagen component (up to 13%). Together with Fig. 2A–C, these findings suggest that combining approaches to reduce SF and adhesion intensities with ECM-targeting strategies, such as collagen-degrading enzymes, may improve CAR-T therapy outcomes. Remarkably, even under conditions of high SF and low adhesion intensities, a 100-fold reduction in collagen not only strongly increases the HF peak—suggesting a condition favorable for complete tumor eradication—but also prolongs hotness within the tumor, potentially lowering the risk of treatment escape. Across our simulations, a 10- to 100-fold reduction in collagen resulted in a 50–200% increase in HF peak and a 49–179% improvement in sustained HF compared with the baseline tumor.

It is possible that metabolic competition between tumor-infiltrating lymphocytes (TILs) and highly metabolic cancer cells can affect tumor hotness. To test this, we examined how cancer metabolism—specifically oxygen and nutrient uptake and carbon dioxide production—affects T cell distribution and HF. Increasing cancer metabolism sharply reduced intratumoral T cell accumulation and HF, with the strongest effects at the highest rates (Fig. 2E). 10- and 100-fold increases in cancer cell metabolism decreased HF by 10% and 86%, respectively. In contrast, reducing metabolism had little effect on the HF peak but substantially prolonged HF duration, allowing T cells to persist even after no new cells were arriving to the tumor.

Strikingly, when both collagen density and cancer metabolism were low, T cells not only persisted but also proliferated and reached a homeostatic equilibrium within the tumor, maintaining a stable population of free T cells over time (Figs. 2D,E). Such conditions could reduce the need for repeated or high-dose CAR-T infusions, which are associated with greater toxicity. While cancer metabolism is variable and difficult to control directly—and also contributes to tumor growth (Fig. 3E)—these results suggest that CAR-T cells engineered for metabolic resistance may be more effective against highly consumptive tumors. Indeed, because aggressive metabolism also imposes growth constraints on cancer cells, metabolically resilient CAR-Ts could exploit this vulnerability and achieve durable tumor control. Establishing equilibrium conditions, especially under low-collagen and low-metabolism states, therefore holds promise for sustaining long-term CAR-T efficacy in tumors.

**Figure 3.**
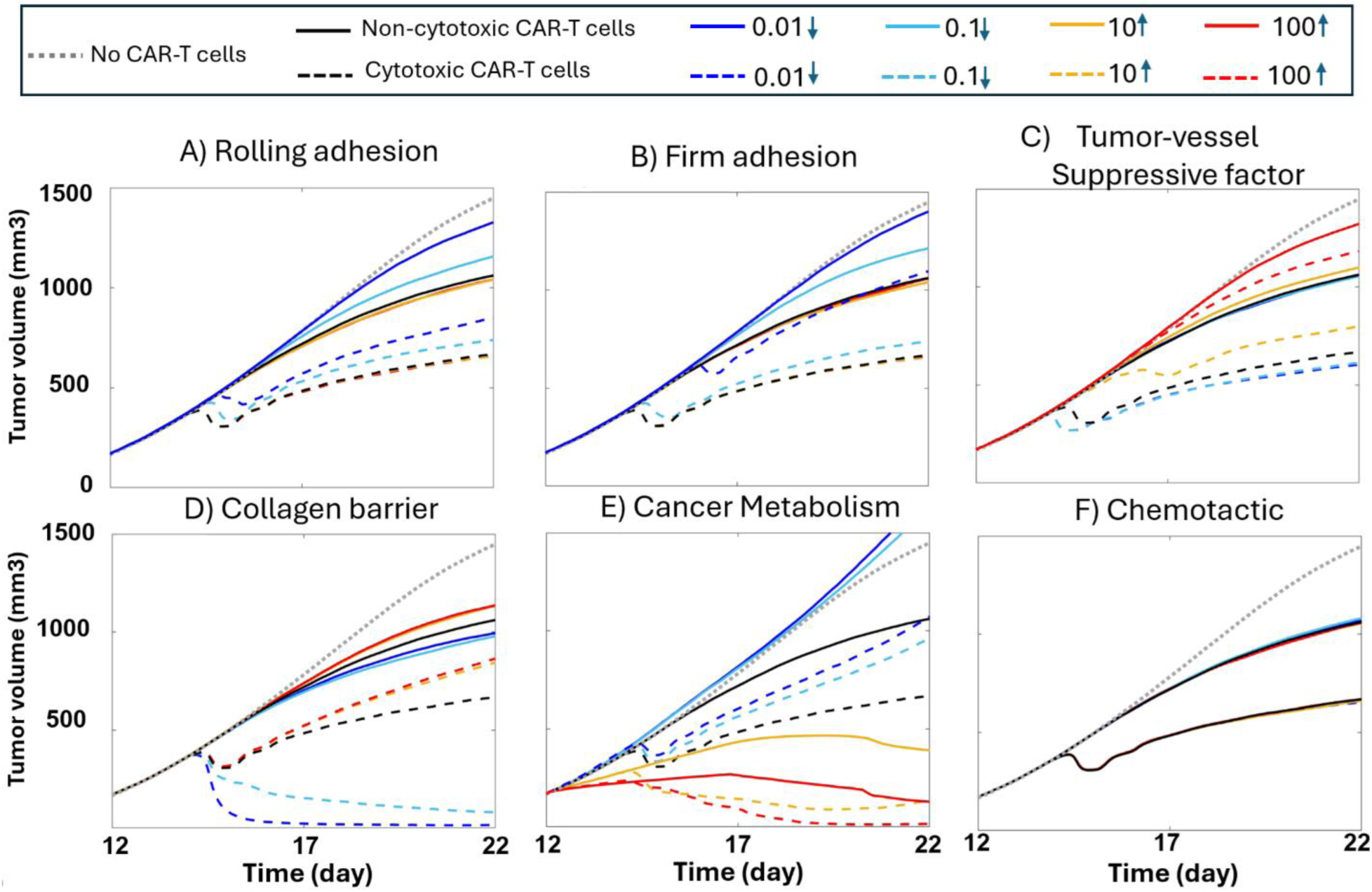
T cell trafficking and cancer metabolism affect CAR-T cell killing of tumors. Solid lines are simulations with non-cytotoxic CAR-T cells; dashed lines are simulations with cytotoxic CAR-T cells; Dotted gray line represents the tumor without T cells (immune-desert condition). A) Tumor growth curves in tumors with different rolling adhesion rate constants. B) Tumor growth curves with different firm adhesion rates, C) Tumor growth curves with different SF levels of tumor vessels, D) Tumor growth curves with different levels of collagen, E) Tumor growth curves with different levels of cancer metabolism rates, F) Tumor growth curves with different levels of chemotactic intensity.

The final extravascular factor influencing T cell–TME interactions is the intensity of chemotactic migration toward tumor-derived chemokines. As shown in Fig. 2F, HF is relatively insensitive to this parameter, as T cells primarily traffic through the pervasive tumor vasculature, making chemotaxis a relatively minor contributor to their intratumoral distribution.

### Intratumoral CAR-T distribution affects therapeutic response

Next, we investigated how the variations in HF translate to tumor growth reduction by allowing the CAR-T cells to kill cancer cells. The tumor growth profiles under different intravascular and extravascular barrier scenarios (Fig. 2) are shown in Fig. 3 for both non-cytotoxic (solid lines) and cytotoxic (dashed lines) CAR-Ts. Simulating non-cytotoxic CAR-T allows us to see how crowding and metabolic processes affect tumor growth, independent of cell killing. Compared to immune-desert tumors without any T cells (dotted line), non-cytotoxic CAR-T cells modestly reduce tumor growth, largely through spatial and metabolic competition with cancer cells. Among intravascular factors, variation in SF had the strongest effect on CAR-T outcomes, with fluctuations up to 91%, whereas firm and rolling adhesion intensities caused smaller fluctuations of ∼58% and ∼38%, respectively. Among extravascular barriers, collagen and metabolism had the greatest impact. Reducing collagen concentration enabled complete tumor eradication by enhancing CAR-T infiltration into both proliferative and quiescent regions, maintaining an intratumor population until all cancer cells were cleared. Although cancer cells could, in principle, refill the spaces vacated by dying cells, their slower migration compared to CAR-Ts allowed the immune cells to eliminate them first. Conversely, increased collagen severely limited CAR-T access and impaired outcomes. Cancer metabolism also strongly influenced therapy: while high metabolic rates reduced HF (Fig. 2E), they also suppressed tumor growth via nutrient competition among cancer cells (solid red and yellow lines in Fig. 3E). In this context, cytotoxic CAR-Ts completely eradicated tumors with maximum metabolism (red dashed line) and induced remission in highly metabolic tumors with possible escape (yellow dashed line). Paradoxically, lowering cancer metabolism accelerated tumor growth (blue solid lines, Fig. 3E) but also prolonged HF (Fig. 2E), allowing cytotoxic CAR-Ts to eventually control tumor growth (blue dashed lines, Fig. 3E). These findings illustrate CAR-T performance across tumor types, from normoxic to highly hypoxic and necrotic tumors, where competition between cancer and immune cells drives heterogeneous outcomes. Importantly, engineering CAR-Ts with enhanced resistance to low nutrients, hypoxia, and metabolite toxicity may ensure consistent efficacy, particularly against highly metabolic tumors (e.g., outcomes resembling the red dashed curve in Fig. 3E). Finally, consistent with the HF results in Fig. 2F, chemotactic intensity had negligible influence on CAR-T efficacy in vascularized tumors, since vessel-mediated entry dominated over chemokine-driven migration.

### Systemic injection of CAR-T cells provides better tumor control compared with intratumoral injections

Because of clinical failures to achieve sufficient intratumor delivery of CAR-T cells ^79–82^, another emerging strategy is to inject the CAR-T cells directly into the tumor ^82–85^. We thus compared intratumor with systemic intravenous (IV) injection, as well as the combination of these two approaches (Fig. 4). For intratumoral infusion, we simulated two scenarios: injection directly into the tumor interstitium (IT) and injection into a large blood vessel feeding the tumor (intra-arterial, IA). Since the simulations predict that migration through the collagen matrix is the dominant factor in HF and CAR-T treatment outcome, we chose tumors with low (0.01× baseline), medium (baseline) and high collagen levels (100× baseline). The results predict that IT infusion is not sufficient for long-term response, even with low collagen levels, as the tumor starts relapsing after a couple of days post-infusion. The best responses were obtained with the combination of intratumor and systemic administration, while systemic alone was also effective. In the low collagen tumor, combined IA and IV infusions also resulted in sustained tumor regression. Although the combination of IT and IA resulted in early control of tumor growth, it did not persist for more than five days post-infusion. Increasing the collagen barrier from low to normal and high reduces the efficiency of IV administration and its combinations with IT and IA. However, combining IA with IT, compared to IT alone produces better outcomes. Increased collagen level reduces this benefit. The best outcomes were produced by the combination of IV and IT with only a slight relapse in the high level of collagen and a complete cure in low and normal levels of collagen.

**Figure 4.**
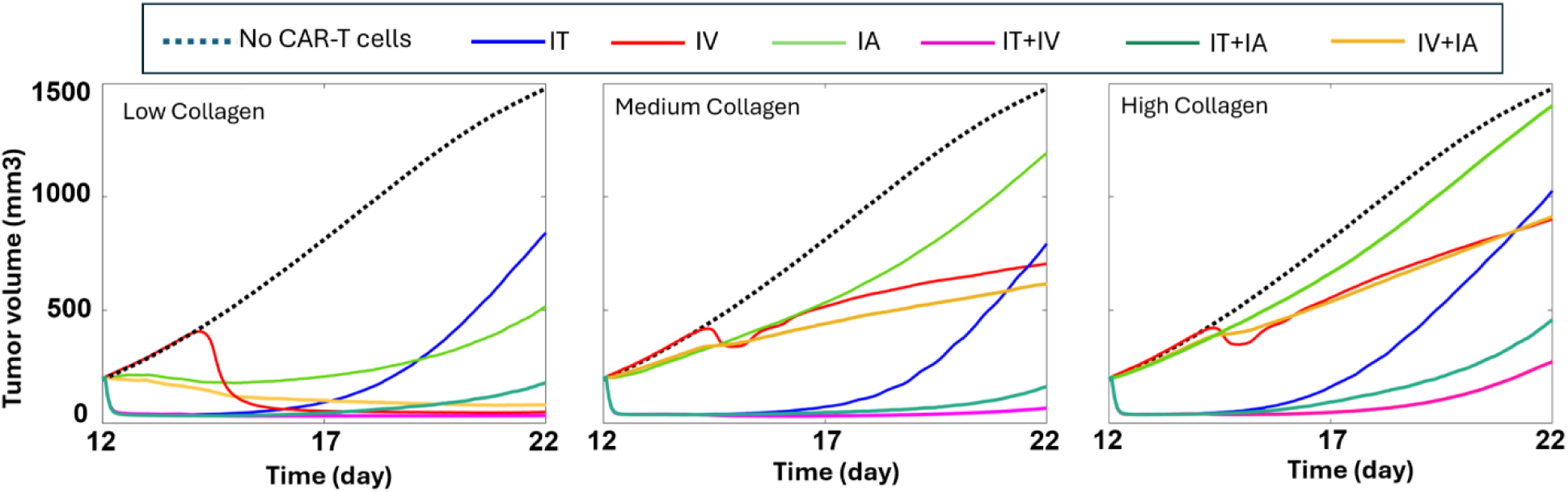
Comparison of cytotoxic CAR-T cell infusions under different collagen levels in the tumor. The middle panel (medium collagen) represents the baseline tumor model, while the left and right panels correspond to 0.01-fold (low) and 100-fold (high) collagen levels in the tumor region, respectively. For intravenous (IV) infusion, CAR-T cells are administered following a Gaussian temporal profile (blue curve in Fig. 1). For intratumoral (IT) and intra-arterial (IA) infusions, CAR-T cells are delivered as short pulse inputs. To ensure a fair comparison across infusion routes, the total number of infused CAR-T cells was normalized by matching the area under the concentration–time curve (AUC) of each administration profile. Tumor volume represents the total tumor mass, including both viable and necrotic regions.

## Discussion

We developed a computational model of the TME to simulate T cell behavior and CAR-T efficiency. The model captures critical mechanisms controlling CAR-T efficacy by incorporating signaling barriers that hinder T cell activity and examining how modifying the TME can enhance T cell delivery and homing in tumors, thereby increasing tumor hotness and the chance of cure. Using a validated 3D vascularized tumor framework and general T cell dynamics in the TME, and by recapitulating morphological and physiological patterns of T cells, we showed that among intravascular factors, tumor-induced vascular suppressive factors have the greatest impact on infiltration, followed by firm adhesion and then rolling adhesion. Increasing SF and reducing firm or rolling adhesion significantly decreases HF; however, improving HF by reducing SF or enhancing adhesion is ultimately constrained by physical barriers from dense collagen and by metabolic competition from cancer cells. Thus, therapies targeting adhesion or SF are most effective when combined with ECM-modifying strategies such as collagen degradation. Importantly, engineering CAR-Ts with enhanced resistance to low nutrients, hypoxia, and metabolite toxicity may ensure consistent efficacy, particularly against highly metabolic tumors. As long as vessels conduct T cells into tumors, tumor-induced chemotaxis has a negligible effect on infiltration and TH, as migration distances are minimized in well-vascularized tumors.

The model further shows that infusion strategy strongly influences short- and long-term CAR-T responses: intratumoral infusion achieves rapid reductions across tumors with different collagen levels but lacks durability, as tumors resume growth within days. In contrast, systemic infusion, leveraging distributed tumor vasculature, provides less immediate control but sustains longer responses—achieving complete cure in low-collagen tumors and partial control in medium to high-collagen tumors. A combined systemic plus intratumoral approach yields the best outcomes, with complete cures in low- to medium-collagen tumors and durable control even under high collagen. Infusion into the tumor interstitium and into the tumor feeding vasculature might further improve outcomes in low-collagen tumors, although performance declines significantly in collagen-rich environments.

In this study, we assumed that all cancer cells originate from a single colony but exhibit different proliferation and migration behaviors depending on their spatiotemporal energy and vitality, determined by local conditions in the TME. Our model shows that cytotoxic CAR-T–mediated killing of cancer cells in certain regions does not necessarily indicate therapeutic progress. When T cells eliminate metabolically active tumor cells in dense regions—typically near functional vessels—they may inadvertently create space and reduce resource competition, thereby allowing highly proliferative cancer cells to expand more rapidly. This effect can also compromise T-cell function, as the highly proliferative cancer cells subsequently starve the T cells. These dynamics explain the fluctuations in tumor volume in the simulations with cytotoxic CAR-T cells (see Fig. 5), particularly in the baseline tumor characterized by high collagen levels, where T cells are closer to vessels and consequently promote local tumor regrowth following the initial reductions.

**Figure 5.**
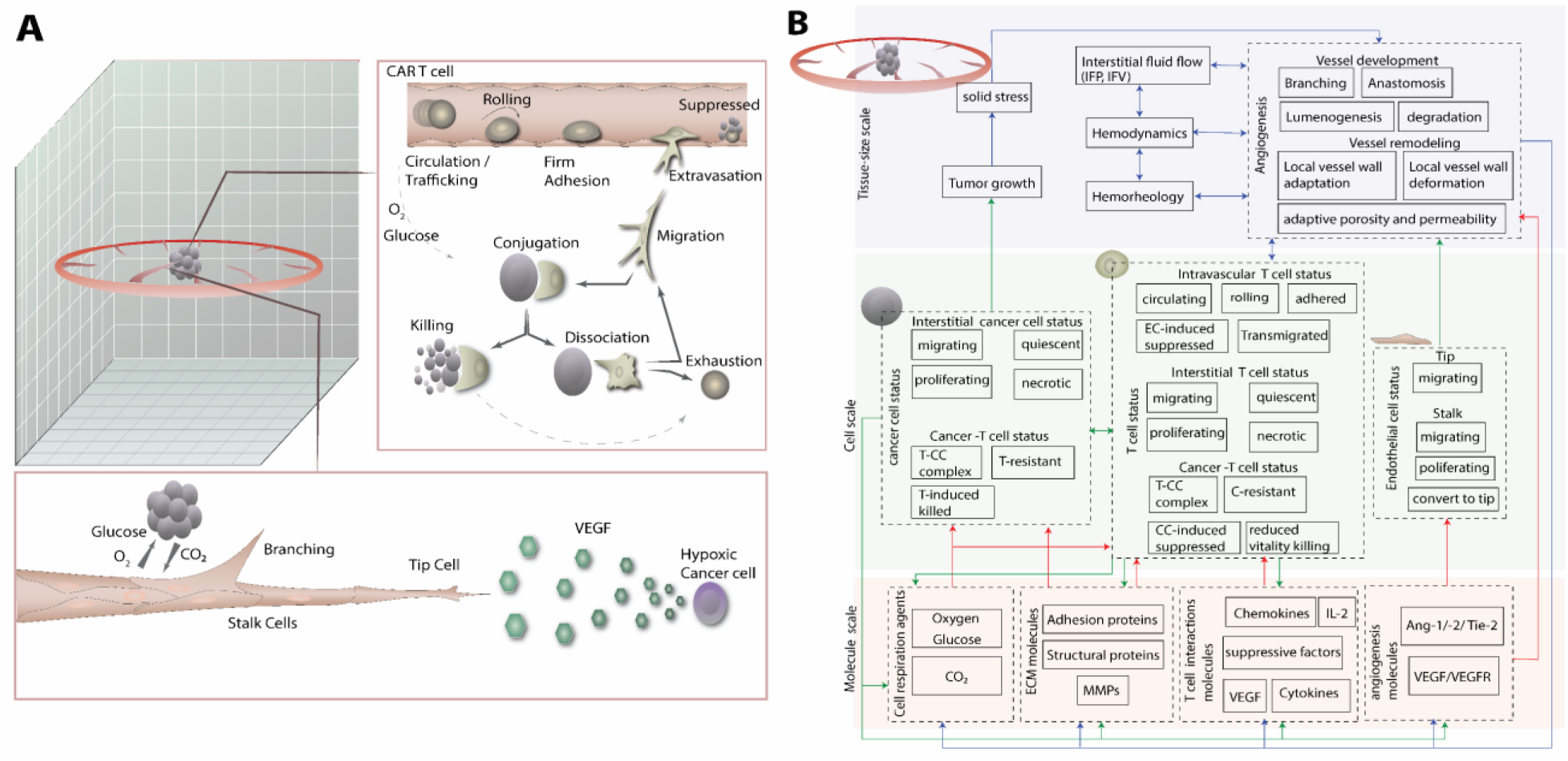
Hybrid multiscale model of T cell interactions within the tumor microenvironment (TME) of a solid vascularized tumor. (A) Schematic of the 3D computational domain, including tumor tissue, angiogenic vessels, and T cells. Circulating T cells can undergo rolling followed by firm adhesion—or directly establish firm adhesion—on endothelial cells (ECs), facilitated by spatially-resolved, tumor-enhanced lymphocyte adhesion molecules but suppressed by tumor-induced inhibitory factors. After adhesion, T cells may extravasate into the interstitium, where they migrate along chemokine gradients toward the tumor. Upon reaching tumor cells, T cells can form conjugates that either detach, kill tumor cells, or become exhausted. Repeated killing cycles can also lead to T cell exhaustion (dashed line). Vessels are dynamic elements that continuously branch and expand their vascular networks in response to local tissue demand and blood flow. (B) Computational framework of the model spanning molecular, cellular, and tissue scales. Spatiotemporal distributions of molecular agents and cellular properties are computed within the continuous domain using the finite difference method (FDM). These outputs determine cellular phenotypes and dynamics, which collectively shape tissue-scale patterns. At the tissue scale, a hybrid continuous–discrete approach integrates discrete agents (cancer, CAR-T and endothelial cells) with continuous fields (hemodynamics and interstitial fluid flow), enabling simultaneous modeling of cell-level behaviors within the tissue microenvironment.

**Figure 6.**
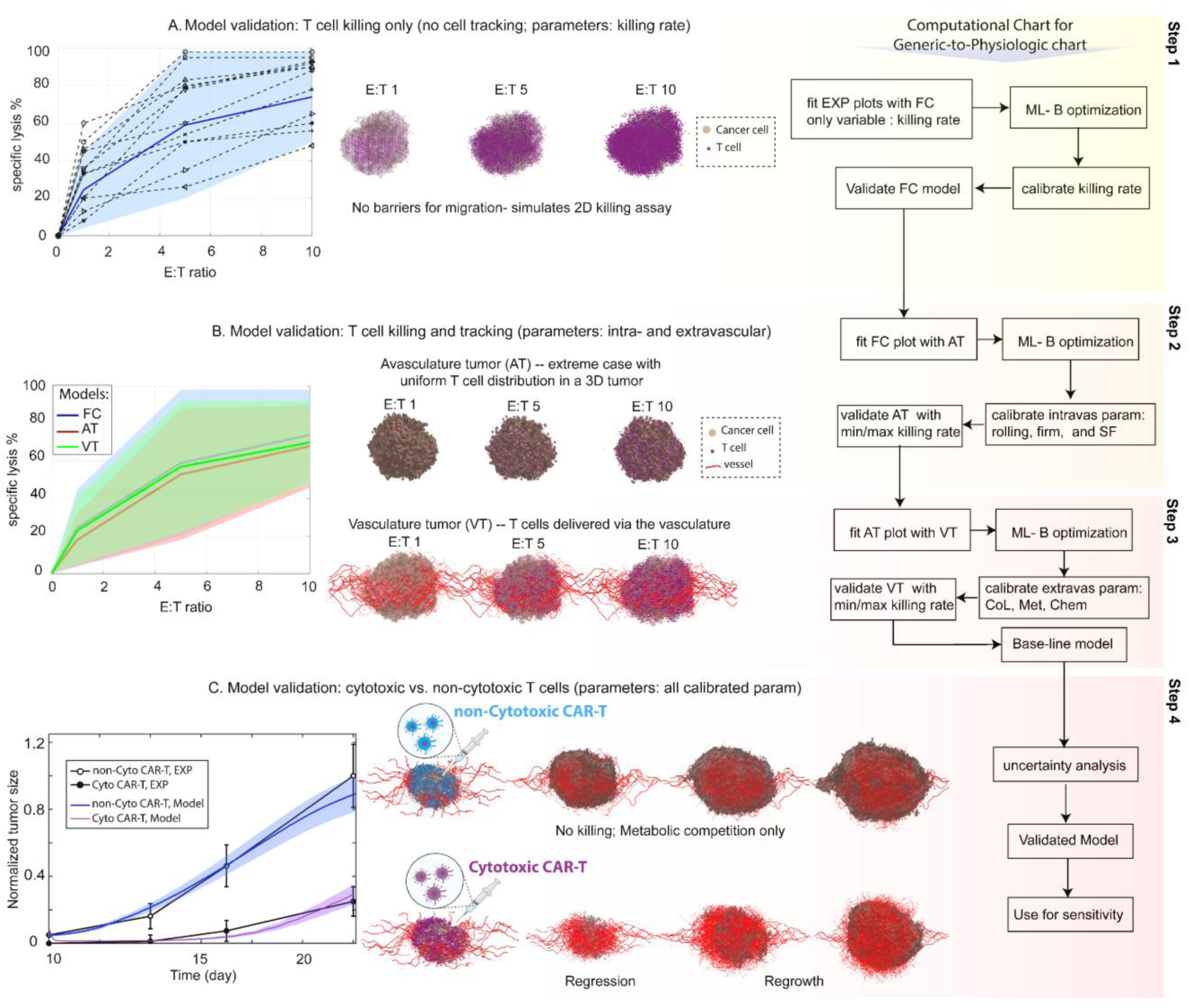
Multi-step validation of model for generic-to-physiology model. A) T cell killing only to calibrate killing rate of CAR-T cells using free cell condition (without 3D barriers for migration and nutrient limitations), mimicking in vitro 2D culture conditions ^73^. The shaded regions represent the range of tumor lysis corresponding to different CAR-T killing rates for each E:T ratio (step 1). B) T cell killing and tracking in 3D avascular (step 2) and vascular tumors (step 3) versus the free-cell model. C) cytotoxic vs non-cytotoxic CAR-T in long-term validation of tumor response to CAR-T across various solid tumors ^74^.

In summary, our computational model demonstrates that CAR-T efficacy is primarily constrained by extravascular barriers such as dense collagen and metabolic competition in the tumor microenvironment, with intravascular factors like tumor-induced vascular suppression and impaired adhesion also playing significant roles; alleviating these through combined therapies, including adhesion enhancement and collagen degradation, can markedly improve T cell infiltration and tumor hotness. Treatment outcomes are further influenced by infusion strategies, where systemic delivery yields sustained responses and superior long-term control compared to intratumoral infusion’s rapid but transient effects, while a hybrid approach combining both methods achieves the best results, including complete cures in low- to medium-collagen tumors and durable control in high-collagen environments—though CAR-T-mediated killing may paradoxically promote tumor regrowth by reducing resource competition. Once validated through preclinical and clinical studies, these insights could inform optimized strategies for enhancing CAR-T therapies across diverse tumor types.

## Acknowledgments

We thank Dr. Timothy P. Padera for helpful suggestions and edits to the manuscript. Lance Munn’s research is supported by NIH R01CA247441, U01CA261842 and R01CA284603.

## Methods

### Generic computational model

Details of the mathematical model can be found in the Supplementary Material file. Briefly, our multi-scale model domain represents a 3D region of TME and considers cancer cells, endothelial cells and T cell dynamics on discrete matrices. The cells can communicate in a continuous field of TME agents. In this field, we calculate continuous gradients of oxygen, nutrients (glucose), and waste metabolites (carbon dioxide), VEGF, ECM proteins (collagen and fibronectin) and matrix metalloproteinases (MMPs), Angiopoietins- 1 and 2 (ang-1 and -2), immune suppressive factors (SF) and pro-inflammatory cytokines, and chemokines. SFs are secreted by tumors and act on vascular endothelial cells to suppress rolling and firm adhered T cells ^9,54–56^. In contrast, pro-inflammatory cytokines promote the upregulation of adhesion molecules on endothelial cells, thereby enhancing T cell rolling and firm adhesion ^57–60^. Additionally, tumor-derived chemokines establish chemotactic gradients within the interstitium that guide T cell migration toward the tumor tissue ^61–64^.

Angiogenic blood vessels initiate from an idealized circular “mother vessel” that surrounds the tumor at the mid-plane (Figure 5.A). Angiogenic sprouts migrate from the mother vessel in a biased random walk toward sources of VEGF and haptotactic factors. Sprout extension requires endothelial cell proliferation.

At each time step, cancer and T cells are assigned a vitality, which is determined by local oxygen, glucose, and carbon dioxide concentrations; cell viability and proliferation depend on this vitality score. Proliferation also depends on the local availability of space, so decreases with high cell density. Cancer cells migrate with a random walk, biased toward oxygen and nutrient gradients, implicitly causing migration towards vessels (vessel cooption). Cancer cell migration is also biased against directions where cell density is higher ^65–67^. Transport of oxygen, nutrients and drugs occurs by convection within the vascular network, and by diffusion and advection through the extravascular tissue. Similarly, fluid and metabolites can enter blood vessels via convection if the local pressure gradient is appropriate. Low oxygen levels trigger the production of VEGF by cancer cells, and VEGF enhances endothelial proliferation and guides the extension of angiogenic vessels. The blood vessels also respond to fluid forces, increasing their diameter in response to higher shear stress ^48,68–70^.

Endothelial permeability is increased by VEGF and the ratio of Ang-1/Ang-2, so transvascular diffusion of plasma and nutrients is increased in hypoxic regions. In addition to limiting cancer cell proliferation and migration, increased cell density can mechanically compress angiogenic blood vessels, decreasing perfusion ^65,71^.

T cells transported via blood flow can interact with the vessel wall through lymphocyte adhesion in response to cancer-derived cytokines. They first transition to a rolling adhesion state, then to firm adhesion, after which they can either be suppressed by tumor-derived vascular inhibitory factors or transmigrate across the vessel wall into the interstitial space. There is also the possibility that the T cells can directly establish firm adhesion to endothelial cells. After transmigration, T cells migrate within the interstitium via a combination of random walk, haptotaxis along ECM adhesion proteins and collagen gradients, and chemotaxis toward positive gradients of tumor-released chemokines ^72^. In this 3D space, cancer cells and T cells compete for physical space, oxygen, and nutrients, mutually constraining each other’s expansion and function. To model this competition, we introduced parameters representing overall cell density as well as cancer cell metabolic activity, referred to as the “cancer metabolism” factor.

#### Computational domain

a 10 ×10 ×10 mm cuboid with 201 × 201 × 201 lattice nodes was selected for the computational domain. The hybrid continuous-discrete (HCD) method defined in our previous TME model ^65^ was applied to solve the mathematical equations of the model. The schematic of the TME computational domain with its different scales, including tissue, cellular, and molecular scales is shown in Fig 5.

#### Computational method

In the HCD computational approach, we divided the model equations into continuous and discrete parts. In the continuous part of the model, the governing equations are numerically solved by an appropriate finite difference method (FDM) on the three-dimensional mesh of the TME cubic domain. The continuous part of the model includes the equations of spatiotemporal distributions of the molecular scale, the cellular vitality and energy equations of the cellular scale, the spatiotemporal distributions of biomechanical factors, vessel growth and remodeling. The discrete part of the model includes cellular density equations defined for cellular dynamics of tumor and tip endothelial cells, and T cells ^65^. Here, the model is discretized on three distinct lattices with the same grids as the finite difference mesh applied to the continuous parts. The computational flowchart is shown in Fig. 5 to illustrate the relationships between different scales of the model and the different computational methods for different scales.

As shown in Fig. 5B, the computational steps after setting the initial and boundary conditions are: 1. Update molecular agents on the finite difference mesh within the same 3D domain as the TME domain (Fig.5A) [O_2_, glucose, and CO_2_ fields, ECM proteins and MMP fields, cytokines and chemokine fields, SF field, VEGF and the VEGF receptor (VEGFR-2) fields, Ang-1, Ang-2 and their common receptor (Tie-2) fields]; 2. Update cellular features on the finite difference mesh [cellular vitality for cancer and T cells, probability of branching of vessels for stalk-to-tip (Eq. 21 in S1 Text), update phenotypes of cancer, T-cells and endothelial cells]; 3. Update the tissue scale on the finite difference mesh for hemodynamics, interstitial fluid flow, tumor-induced solid stress, and vessel growth and remodeling variables; 4. Update the tissue scale on the lattice of tumor cells for tumor growth; 5. Update the tissue scale on the lattice of ECs for angiogenesis; 6. Update the lattice of the intravascular T-cells for status, including circulating (free T cells), rolling T cells, firmly adhered T cells, suppressed T cells, and extravasated T cells 7. Update the lattice of the extravasated T cells using the interstitial T cell density equation with biased random walk with haptotactic and chemotactic mechanisms (Eq. 27 in Supplementary Material) 8. Update the lattice of interstitial T cells using interactions with cancer cells, including conjugation, dissociation, killing (when CAR-Ts are in the cytotoxic phase), pre-cancer-killing immediate exhaustion, post-cancer-killing delayed exhaustion, 9. Update the molecular and then cellular scales based on the updated tissue scale information.

For the simulations in this study, a tumor was “grown” to a size of ∼200 mm^3^, and this domain was then used for the initial domain state for all simulations. Note that angiogenesis mechanisms continue to be active during tumor growth and CAR-T treatment, so the vasculature is dynamic in the simulations that follow.

### Physiological model

To develop a physiological model using the generic framework, we established a novel multi-layer “generic-to-physiologic” modeling approach that integrates successive levels of biological complexity: (A) a free-cell model (FC) representing cancer–T cell interactions in an unbounded domain without nutrient or spatial limitations, (B) a 3D model incorporating tumor and vascular structures, and (C) a long-term response analysis for validation. In this framework, all model parameters listed in Table A (Supplementary Material) were fixed except for six key parameters related to intravascular and extravascular barriers affecting T cell trafficking and CAR-T cytotoxicity. The *intravascular parameters* included: (a) T cell rolling adhesion efficiency along the vessel wall, (b) T cell firm adhesion efficiency to the endothelium, and (c) the intensity of tumor-induced suppressive signaling on endothelial cells that modulates adhered T cell activity. The *extravascular parameters* included: (d) collagen density, (e) the intensity of metabolic competition between cancer and T cells, and (f) T cell chemotactic strength.

As shown in Fig. 6 (computational chart of the generic-to-physiologic model) we performed the following steps to calibrate the model: **Step 1:** The goal was to calibrate the CAR-T killing rate using experimental data from Mishra, et al. ^73^ under various effector-to-target (E:T) ratios (0, 1, 5, 10) after 16 h. We employed the FC model, which excludes spatial and nutrient constraints, mimicking a 2D co-culture of CAR-T and cancer cells. Using a Bayesian Machine Learning (BML) optimization algorithm, the model was fitted to the experimental outcomes to obtain an averaged killing rate consistent with reported CAR-T activity in solid tumors. **Step 2:** Next, we calibrated the three *intravascular* parameters—rolling, firm adhesion, and suppressive intensity (a–c)—using an Avascular 3D tumor (AT) model that introduces spatial barriers such as metabolic competition and collagen density. The same E:T ratios were applied, and BML was again used to fit the FC model’s killing curve (from Step 1) to the AT model results. After calibration, we compared killing outcomes for the minimum and maximum fitted rates, showing that the averaged AT curve reflects the attenuated CAR-T cytotoxicity in 3D due to physical and metabolic barriers (red vs. blue line, Fig. 6B). **Step 3:** The next goal was to calibrate the three *extravascular* parameters (d–f) using the 3D vascularized tumor (VT) model. BML optimization was performed by fitting the AT model killing curve (from Step 2) to the VT results using the same E:T ratios (1, 5, 10). The calibrated VT model was then simulated across minimum and maximum values, and the averaged response remained within the valid range—positioned between FC and AT curves—reflecting the positive role of vasculature in enhancing CAR-T access and function. This model was selected as the baseline physiological model. **Step 4:** Finally, we validated the baseline physiological model by simulating long-term CAR-T–tumor interactions under both cytotoxic and non-cytotoxic CAR-T conditions. The non-cytotoxic phase captured only metabolic competition in a confined domain, whereas the cytotoxic phase included both metabolic competition and direct tumor cell killing. As shown in Fig. 6C, the baseline model accurately reproduced experimental long-term CAR-T responses across multiple solid tumor types^74^, demonstrating strong agreement between model predictions and observed CAR-T kinetics in both cytotoxic and non-cytotoxic regimes. Due to the stochastic nature of cellular interactions, multiple outcomes can emerge from the same parameter set, represented as shaded regions. These regions illustrate the uncertainty of the physiological model predictions, while the averaged results show good agreement with the experimental data.

The localization of T cells within different interstitial regions of the TME—including within normal vessels distant from the tumor, within peritumoral vessels, and within intratumoral vessels—is shown in the Supplementary Material (Figs. S2 and S3), consistent with experimental observations.

